# Evaluating Spike Antigenicity across Endemic Human Coronavirus Models using Flow Virometry

**DOI:** 10.64898/2026.05.28.728498

**Authors:** Jonathan Burnie, Caroline Ouano, Victor Luo, Christian K.O. Dzuvor, Taiya Miller, Giancarlo Ospina, Nikhila S. Tanneti, Li Hui Tan, Donald J. Hamel, Catherine Hammond, Hannah Matthews, Lamount R. Evanson, Jeswin Joseph, Sydney P. Moak, Phyllis Kanki, Noam A. Cohen, Susan R. Weiss, Kizzmekia S. Corbett-Helaire

## Abstract

While SARS-CoV-2 research has advanced rapidly since COVID-19, endemic human coronaviruses (HCoVs) remain comparatively understudied. Tools to phenotype spike (S), the primary antigenic target on coronaviruses, at the single-virion level could improve vaccine design by capturing variation in epitope availability and spike abundance. Here, we establish a calibrated flow virometry (FV) platform to quantify S antigenicity on native endemic (HCoV-229E, HCoV-OC43) and epidemic (SARS-CoV-2) coronaviruses directly in cell culture supernatants. FV revealed cell line–dependent differences in S antigenicity, including receptor-induced changes in epitope accessibility. Comparison of virion-associated S with recombinant stabilized S by ELISA and biolayer interferometry showed consistent binding for HCoV-OC43, MERS-CoV, and SARS-CoV-2, but differences for HCoV-229E, with FV resolving heterogeneity not captured by bulk assays. Finally, FV showed that HCoV-229E from patient-derived air–liquid interface cultures exhibited reduced antibody binding and distinct S antigenicity compared to cell line–derived virions. Together, these findings establish FV as a platform for single-virion analysis of HCoV antigenicity.

## INTRODUCTION

In recent years, improvements in flow cytometry instrumentation have enabled flow virometry (FV) to emerge as an increasingly feasible, high-throughput method for studying individual virions (reviewed in^1– 4^). FV addresses an important need for approaches that can resolve virus-to-virus variability within heterogeneous populations at the single-particle level. While FV methods for enumerating fluorescently labeled marine viruses have been established for decades^5,6^, the utility of the technique has since expanded to include the analysis of specific viral components—such as surface proteins^7–11^ and capsids^12–15^—and to permit viral sorting^16–18^. The high throughput capacity and rigor of FV have also positioned it as a valuable tool for vaccine quality control^19,20^, with prospective applications in viral diagnostics^21–23^. Historically, FV approaches necessitated fluorescently tagged proteins or dyes to permit detection of viruses on conventional cytometers since most viruses fall below the detection threshold for light scatter, rendering them indistinguishable from background instrument noise^12–14,24,25^. To overcome this limitation, magnetic nanoparticles or bead-based approaches on conventional cytometers^23,26–31^, as well as specialized instrumentation^16,21^, have been employed to enable detection of viruses and study surface proteins on HIV, dengue virus (DENV), influenza, and epidemic human coronaviruses (CoVs), including MERS-CoV and SARS-CoV-2. However, many newer cytometers can readily detect single virions through light scattering alone^9,32–35^, permitting staining of viral surface proteins directly on viruses in cell culture supernatants without the need for magnetic beads, fluorescent tags, ultracentrifugation, or other auxiliary methods^7,9,10,18,32–34,36–38^. Removing these additional requirements streamlines experimental workflows and reduces both costs and potential opportunities for biased data interpretation^1^.

Although CoVs have been a major research focus since the emergence of SARS-CoV-2^39,40^, FV has not been widely applied to characterize the antigenicity of CoV spike (S) proteins. This is especially true for the endemic human coronaviruses (HCoVs; 229E, OC43, NL63, HKU1), which account for 10–30% of upper respiratory tract infections in adults^41^. Beyond their high prevalence, CoVs can exacerbate disease severity in vulnerable populations and in the context of underlying comorbidities (e.g., cardiovascular and metabolic disease)^42,43^. Their endemic circulation also makes HCoVs a valuable model for studying S antigenicity in the context of vaccine design and coronavirus immune responses. Given their endemic circulation, pandemic potential^44^, and ongoing zoonotic risk^45,46^, advancing our understanding of CoV biology remains a critical priority.

A key challenge in studying CoVs is their pleiomorphic nature^47,48^, producing heterogeneous virion populations that vary in size, spike density, and composition, features that can directly influence antigenicity and antibody accessibility. This heterogeneity may be further amplified by the presence of defective interfering particles^49,50^ with differential spike incorporation and antigenic profiles, properties that are obscured by bulk assays. Accordingly, single-particle approaches are essential for resolving this diversity and informing the development of broadly protective, universal coronavirus vaccines.

To this end, defining the native antigenicity of the S protein is central to guiding vaccine development strategies for broad CoV protection since S is the key viral determinant of entry and a major antibody (Ab) target^51,52^. Within S, the receptor-binding domain (RBD) mediates host receptor engagement and is the primary target of many potent neutralizing antibodies, but is also highly variable across CoVs^53,54^. In contrast, the S2 subunit, which drives membrane fusion, is more conserved and represents an important target for cross-reactive and broadly protective antibody responses^53,54^. Together, these features highlight the importance of resolving antigenic determinants across both variable and conserved regions of S when evaluating CoV immunity and vaccine design.

While single-particle FV studies have begun to emerge characterizing SARS-CoV-2^55–59^, similar antigenic profiling studies of endemic HCoVs have not yet been established. Moreover, most studies of CoV S antigenicity rely on recombinant S proteins stabilized in the prefusion state with two proline mutations^60,61^ (S-2P), as used in many vaccine platforms^62,63^. However, such bulk measurements average signal across large populations of molecules and do not capture the heterogeneity in spike presentation that exists across individual virions. FV provides a unique means to evaluate S antigenicity directly on native virions at the single-particle level, enabling quantitative comparison of antibody binding to recombinant versus virion-associated S and clarifying how faithfully stabilized S constructs recapitulate authentic viral antigenic display. By resolving antigenicity on a per-particle basis, FV can distinguish differences arising from variation in spike density, conformation, or particle integrity that are not accessible through ensemble approaches. However, systematic antigenic profiling of endemic HCoVs at the single-particle level remains limited, and direct comparisons of antibody binding measurements between FV and established bulk assays such as ELISA across multiple coronaviruses are lacking, hindering cross-platform interpretation of spike antigenicity.

Herein, we establish methods to antigenically profile endemic HCoVs (229E and OC43) and epidemic CoVs (MERS and SARS-2) by FV directly in cell culture supernatants, without the need for purification or immunomagnetic enrichment. Using FV, we uncover substantial viral heterogeneity across endemic HCoVs propagated in different human cell lines, as well as differences in spike antigenicity between pseudotyped and replication-competent viruses. Comparing FV with ELISA and biolayer interferometry (BLI), we observe largely consistent antibody binding patterns across S2-directed antibodies for beta-CoVs, with FV uniquely resolving virion-level heterogeneity not captured by bulk assays. We further demonstrate that FV can sensitively detect changes in virion staining at both RBD and S2 epitopes when assays are performed in the presence of soluble cellular receptors. Extending this approach to HCoV-229E virions derived from patient-relevant air–liquid interface (ALI) cultures, we observe reduced and more heterogeneous antibody binding, with donor- and cell-dependent differences in spike antigenicity. Taken together, these data show that FV is a powerful and versatile tool for studying viral surface proteins, uncovering heterogeneity in virus preparations, and offering insights into Ab landscapes and conformational states at single-particle resolution.

## RESULTS

### Establishing FV for single-particle evaluation of S antigenicity across HCoVs

While FV has proven to be a powerful tool for studying viral surface proteins^9,11,26,29,36,55,64,65^, it has not been applied to assess the antigenic landscape of HCoV S proteins. To establish an FV-based platform for HCoVs, we propagated HCoV-229E in three permissive human cell lines representing distinct tissue origins — MRC-5 (lung fibroblast), Huh7.5 (liver hepatoma), and HCT-8 (intestinal epithelial)^66–68^— and labeled S using a quantitative indirect FV staining approach with primary anti-S monoclonal Abs (mAbs) and a PE-conjugated secondary Ab (Fig. S1A). This approach enables the use of unconjugated mAbs and allows fluorescence calibration to molecules of equivalent soluble fluorophore (MESF), providing an estimate of labeled proteins per virion^69,70^.

To assess S antigenicity, we used three mAbs described by Xiang et al.^71^ targeting the RBD (C04), S1 (F07), and S2 (F12) domains of HCoV-229E S. When staining MRC-5-derived HCoV-229E, readily detectable labeling was observed across S for all mAbs, with PE MESF values ranging from 50–115 (∼3– 6-fold above background; Fig. 1A–B), with the highest staining detected for the S2 Ab (115 PE MESF). While the RBD mAb exhibited the lowest level of staining among the three antibodies tested (Fig. 1A-B; 50 PE MESF), this was consistent with neutralization, interaction kinetics and KD values reported by Xiang et al.^71^. For MRC-5 derived virions, all three mAbs enabled clear separation of stained virions from background fluorescence (Fig. 1A, top row) which were significantly different from the isotype control (Fig. 1B).

**Figure 1.**
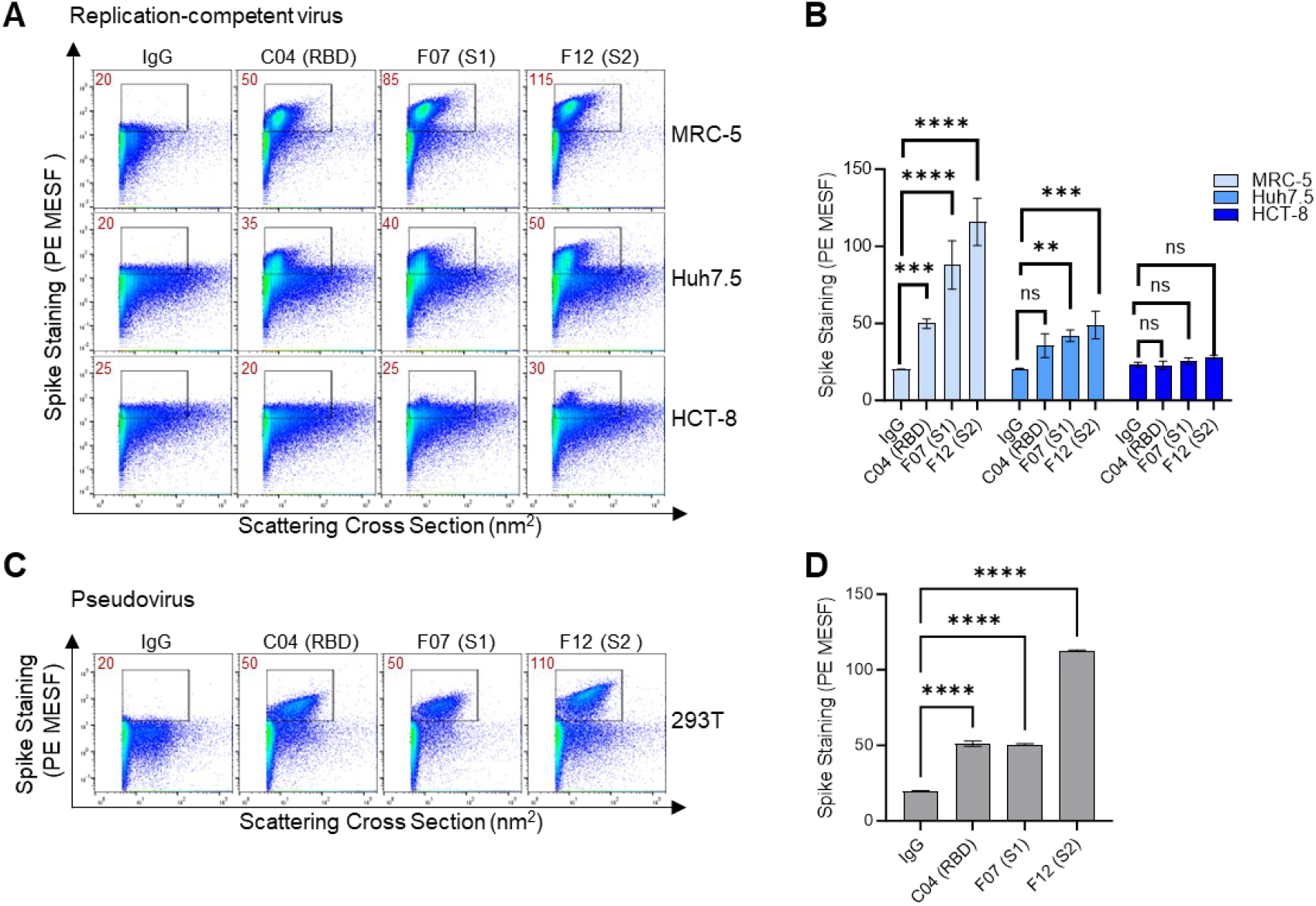
Establishing an FV platform to assess S antigenicity on HCoV-229E. (A) Supernatants from HCoV-229E-infected human cell lines (MRC-5, Huh7.5, and HCT-8; top to bottom rows) were stained using S-specific mAbs targeting the RBD (C04), S1 (F07), or S2 (F12) domains, or an isotype control (IgG), followed by a PE-labeled secondary antibody. Gates for positive staining were set above the isotype control background (>15 PE MESF on the y-axis). Calibrated mean PE fluorescence values (MESF) from gated regions are shown in red. (B) Quantitative comparison of staining results from the gates in (A), with bars representing mean PE MESF ± standard deviation wherein light blue = MRC-5, blue = Huh7.5, and dark blue = HCT-8. Data are representative of at least three independent replicates. Statistical significance was determined using two-way ANOVA with Dunnett’s multiple comparisons test to compare each antibody condition to the IgG control within each virus. Significance thresholds: **p < 0.01, ***p < 0.001, ****p < 0.0001.(C) HCoV-229E pseudoviruses produced by transfection of 293T cells were stained directly in cell culture supernatants using the same antibody panel and staining protocol as in (A). (D) Quantification of staining in (C), with bars indicating mean PE MESF ± standard deviation. See also Figures S1 - S3.

Notably, S antigenicity varied by producer cell line. Compared to MRC-5-derived virions, staining intensity was substantially reduced for viruses generated in Huh7.5 (35–50 MESF) and HCT-8 cells (25 MESF). MRC-5- and Huh7.5-derived virions exhibited comparable infectivity despite these differences, indicating that S antigenicity can vary independently of viral infectivity. In contrast, HCT-8-derived HCoV-229E showed reduced TCID50 values, which may further contribute to its diminished S antigenicity (Fig. S1C).

We next applied this approach to HCoV-OC43 propagated in the same cell lines, as well as A549 (lung epithelial) cells. The CTD-targeting mAb (H501-008)^72^ yielded strong staining across all conditions (∼100–600 MESF; 3–20× above background), whereas the S2-targeting mAb (H501-101)^72^ showed lower signal (40–90 MESF), suggesting reduced epitope accessibility (Fig. S2A) as expected given the immunodominance of CoV S1. Specifically, HCoV-OC43 virions derived from HCT-8 cells exhibited both the highest staining intensity and the greatest heterogeneity, spanning a broad range of side scatter values (5–10,000 nm^2^) and PE MESF (25–40,000; Fig. S2A, C).

Notably, FV resolved multiple distinct particle populations within individual samples, particularly for viruses derived from MRC-5 and Huh7.5 cells (Fig. S1A, panel 4 circles, S2B). While the underlying basis of this heterogeneity—potentially reflecting extracellular vesicles containing spike protein^73,74^ or pleiomorphic virions with variable size and spike abundance^47,48^—is beyond the scope of this study, these data highlight a key advantage of FV to resolve heterogeneity.

Since pseudoviruses (PV) are commonly used in studies assessing how S mAbs affect viral entry ^60,62^, we also tested the same Abs on PV. HCoV-229E PV displayed pronounced staining for all Abs tested (Fig. 1C), particularly for the S2 Ab (F12), with MESF values comparable to those seen with MRC-5 229E reaching 110 ± 0.6 MESF (Fig. 1D). PV populations also showed clear separation from background for all Abs, unlike virions derived from Huh7.5 or HCT-8 cells. In contrast, HCoV-OC43 PV exhibited substantially lower signal than infection-derived virions (Fig. S2D–E), indicating differences in epitope presentation across virus models. These results demonstrate that PV do not uniformly recapitulate native S antigenicity and require empirical validation.

To assess whether the fluorescent signal in FV could be improved, we compared direct staining using a PE-conjugated H501-008 Ab to indirect staining. However, indirect staining yielded 4-fold higher signal (Fig. S3) and thus, was used for all subsequent experiments.

### Detecting changes in S epitope availability when staining in the presence of soluble cellular receptors

CoV S is known to adopt multiple conformational states on virions, including a range of pre- and post-fusion forms^51,75–77^, which influence infectivity, and vaccine or therapeutic antibody design^78,79^. To assess the sensitivity of FV for detecting conformational changes in S, we examined whether FV could resolve receptor-induced alterations in epitope accessibility on intact virions, as previously demonstrated for HIV^80,81^.

To begin, we used pseudotyped HCoV-229E particles due to their broader dynamic range of antibody staining (Fig.1). Virions were pre-incubated with the soluble receptor aminopeptidase N (CD13/APN) for 20 minutes prior to staining with a panel of mAbs targeting distinct S epitopes. This panel consisted of the mAbs used before and S2 antibodies (fp.006^82^, 76E1^83^, 54043-5^84^).

Using this approach, FV resolved epitope-specific changes in mAb binding upon receptor engagement (Fig. 2A–B). No differences were observed in isotype control staining, confirming signal specificity. As expected, RBD-directed staining (C04) was reduced in the presence of APN, consistent with receptor occupancy, although this decrease did not reach statistical significance, likely due to the modest signal seen with mAb C04. Similarly, the S1 antibody F07 showed no measurable change. In contrast, distinct S2-directed mAbs exhibited divergent responses (Fig. 2). The fusion peptide antibodies 76E1 and fp.006 demonstrated increased staining in the presence of APN, suggesting enhanced epitope exposure following receptor engagement. This was in alignment with literature showing that cryptic epitopes within the S2’ site or fusion peptide become more accessible upon receptor engagement^82,83^. Conversely, the S2 apex–targeting mAb 54043-5 showed decreased binding under the same conditions, likely due to its prefusion S2 epitope^84^ becoming less available upon receptor engagement. The F12 mAb, which targets the S2 connector domain, showed no significant change in binding, indicating that distinct epitopes within S2 can be differentially affected by receptor engagement. Applying a similar approach to MERS-CoV pseudoviruses, FV likewise resolved receptor-dependent differences in antibody binding in the presence of DPP4 (Fig. S4), indicating that this strategy is broadly applicable across coronaviruses. Together, these results establish FV as a sensitive platform for detecting subtle, epitope-specific changes in S epitope accessibility on intact virions following receptor engagement.

**Figure 2.**
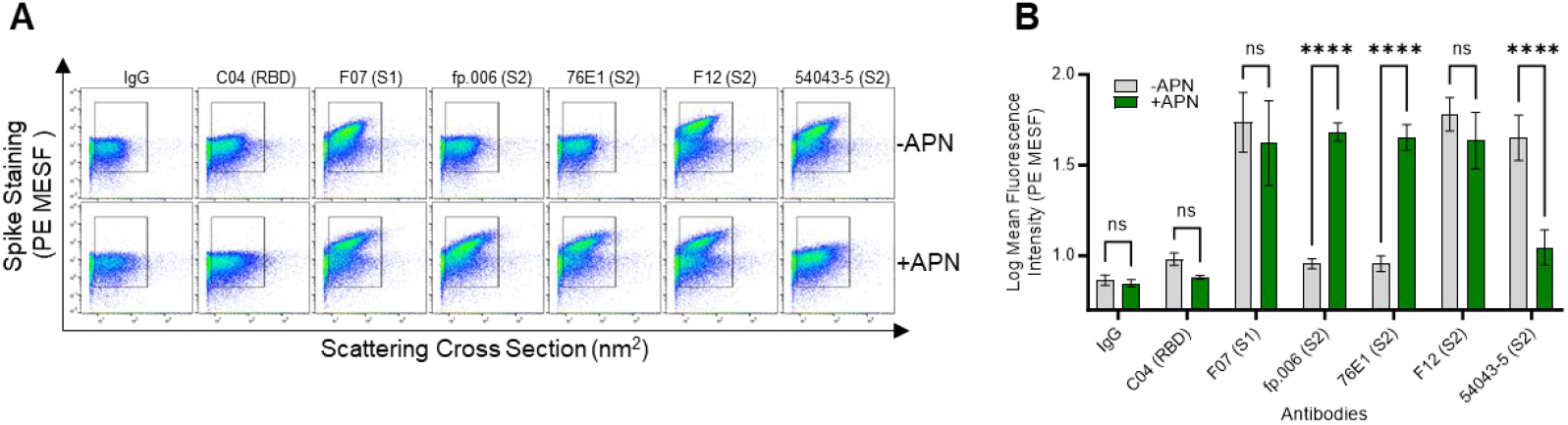
Assessing S epitope availability in the presence of soluble receptor, APN. (A) Flow virometry dot plots showing HCoV-229E pseudovirus staining in the presence (bottom row) or absence (top row) of soluble APN. Gates are drawn on the total virus population to detect modest quantitative changes in staining intensity. (B) Quantification of staining in (A), showing mean PE MESF ± SD from the gated regions of three technical replicates from one experiment, wherein gray = absence of APN and green = presence of APN. Statistical significance was assessed on log10-transformed data using two-way ANOVA with repeated measures and Sidak’s multiple comparisons test (**** = P < 0.0001). Data are representative of three independent experiments. See also Figure S4.

### Comparing binding of cross-reactive S2 mAbs in FV to bulk techniques

Since a major advantage of FV is the ability to visualize antibody binding at the level of individual virions, we sought to compare its performance to a traditional bulk method for assessing antigenicity. To this end, we evaluated a panel of seven cross-reactive S2-targeting mAbs relevant to universal vaccine design (fp.006, 76E1, 54043-5, IgG22^85^, S2P6^86^, and CC40.8^87^) using indirect enzyme-linked immunosorbent assay (ELISA). We tested mAb binding against S-2P^60,61^, a stabilized spike construct commonly used in SARS-CoV-2 vaccine design^62,88,89^ and expanded analysis to a broader range of CoV including HCoV-229E, HCoV-OC43, SARS-CoV-2, and MERS-CoV.

For HCoV-229E S-2P ELISA, fusion peptide mAbs fp.006 and 76E1 showed the highest levels of binding (Fig. 3A). S2P6 showed low levels of binding and all other mAbs did not exceed background binding. Conversely, strong binding was observed across all the mAbs for HCoV-OC43 and SARS-CoV-2 as anticipated (Fig. 3A). Similarly, all mAbs bound well to MERS-CoV S-2P except for CC40.8, consistent with a prior reports^90^ (Fig. 3A). These trends were also largely reflected in biolayer interferometry (BLI; Fig. S5), another commonly used bulk technique for assessing antibody binding to S. To compare these bulk measurements with antibody binding on S from virions, we next performed FV. For this analysis, replication-competent viruses were used in all cases except MERS-CoV, which was evaluated using pseudovirus due to its BSL-3 biosafety requirements.

**Figure 3.**
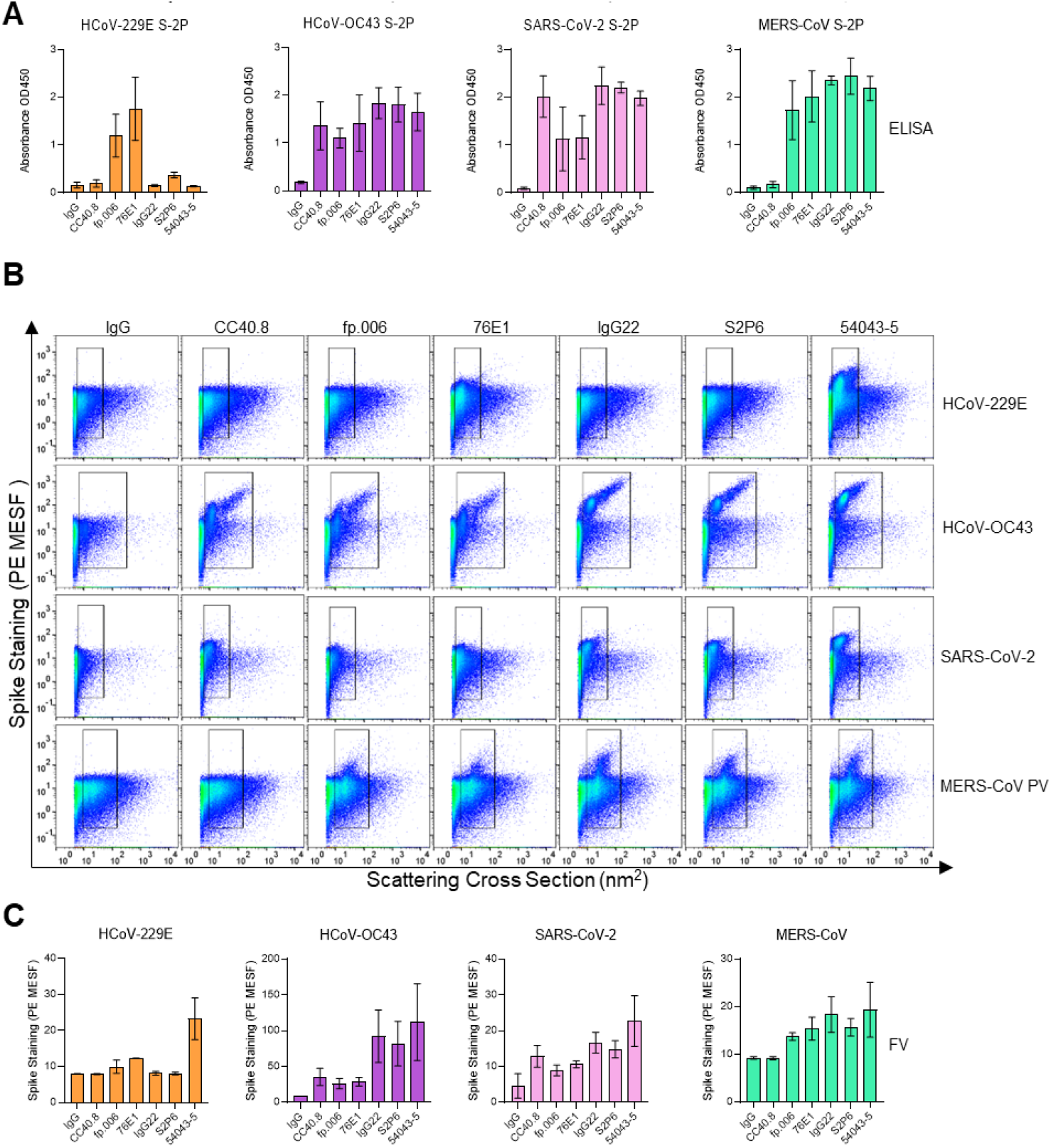
Comparing cross-reactive S2 antibody binding against HCoV in FV and ELISA. (A) Binding of mAbs to recombinant S-2P as determined by ELISA. Data represent at least two independent experiments, each with triplicate wells. Data are representative of two independent experiments with staining performed in duplicate wells. (B) Replication-competent HCoV-229E and HCoV-OC43 grown in MRC-5 cells, HEK293T-derived MERS-CoV pseudoviruses, and SARS-CoV-2 propagated in A549-ACE2 cells, were stained with a panel of S2-specific mAbs and analyzed by flow virometry. Gates represent total antibody-stained virus populations relative to the isotype control (IgG). (C) Quantitative comparison of staining results from the gates in (B). See also Figure S5.

We found that FV revealed notable differences from binding in ELISA. While fp.006, 76E1, and S2P6 all showed binding in ELISA, flow analysis revealed minimal binding for fp.006, no detectable binding for S2P6, and reduced but detectable staining for 76E1. The strongest staining was observed for 54043-5, in contrast to the S-2P ELISA results. Interestingly, for HCoV-229E, two distinct scatter populations were observed with 54043-5 staining within the virus gate (Fig. 3B; green vs. blue scatter population on the left and right, respectively). This was less evident for 76E1, likely due to its lower overall staining. While the biological relevance of these distinct populations will be the subject of future work, their presence highlights a key limitation of bulk techniques such as ELISA, which do not resolve virion heterogeneity at the individual particle level.

For HCoV-OC43, all antibodies exhibited robust staining. Notably, two clearly distinct staining populations were observed across all mAbs, more distinctly defined than those seen for HCoV-229E. For SARS-CoV-2 staining was seen across all the mAbs, similar to ELISA, however, at lower levels for fp.006 and 76E1. Of note, these viruses displayed one, monodisperse population. Finally, MERS-CoV PV displayed mAb binding consistent with ELISA and a single monodisperse virus population. However, only a subset of virions showed spike staining above background for all mAbs tested, in contrast to the clear population shifts observed for HCoV-OC43 and SARS-CoV-2 with strongly staining mAbs such as 54043-5.

Importantly, although FV data summarized as population means showed overall trends largely consistent with ELISA for OC43, SARS-2 and MERS (Fig. 3C), FV dot plots provided a more nuanced visualization of single-virion heterogeneity. Together, these data demonstrate that antibody binding to native virions is not uniform, and that subpopulations of particles differ in epitope accessibility—features that are not captured by bulk ELISA measurements. This further suggests that antibody binding to S-2P may differ from staining for both native virions and PV.

### Assessing S antigenicity of HCoV-229E grown in patient-derived nasal and bronchial cells cultured at the air liquid interface

Since propagation in cell lines can introduce mutations that alter S antigenicity^91,92^, we next used FV to investigate biological differences in S antigenicity in viruses grown in patient-derived air–liquid interface (ALI) cultures (Fig. 4A)^93^, a model with greater physiological relevance. ALI cultures closely mimic the *in vivo* epithelium in cellular composition (ciliated, goblet, and basal cells) and function (barrier integrity, mucociliary activity), providing an optimal system to study HCoVs^94–96^. To this end, we cultured viruses in paired nasal (upper airway) and bronchial (lower airway) cells from four donors for FV staining since both tissue types are sites of CoV replication *in vivo*^95,97^.

**Figure 4.**
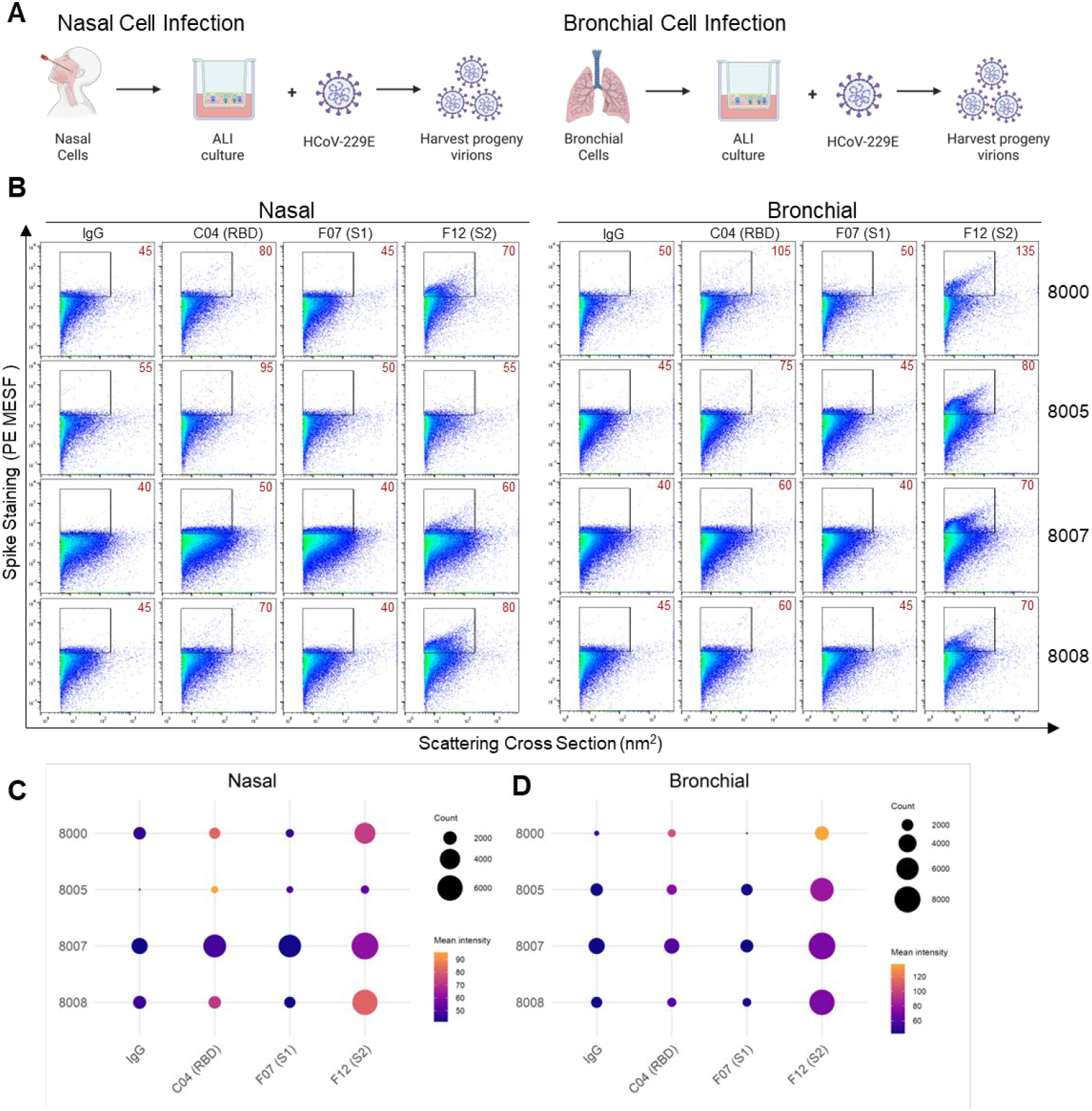
Assessing S antigenicity in HCoV-229E virions produced in primary air liquid interface cell cultures. (A) Schematic depicting propagation of viruses in air liquid interface (ALI) cultures derived from nasal or bronchial cells. (B) HCoV-229E propagated in four different donor cells (8000, 8005, 8007, and 8008 – top to bottom rows) cultured at ALI were stained using S-specific mAbs targeting the RBD (C04), S1 (F07), or S2 (F12) domains, or an isotype control (IgG), followed by a PE-labeled secondary antibody. Gates are set based on the isotype control and extend above background fluorescence (>30 PE MESF). Calibrated mean PE fluorescence values (MESF) from gated regions are shown in red. Results are representative of three technical replicates. (C) Quantification of mean staining fluorescence (PE MESF) in nasal-derived or (D) bronchial-derived virions from (B). Bubble color represents PE MESF (staining intensity) and bubble size corresponds to the number of events (Count) detected for each antibody–virus combination.

While cell line-derived HCoV-229E showed strong staining with all the S mAbs tested (C04, F12, F07; Fig. 1), viruses grown in the ALI model showed notable staining only with the S2-specific mAb F12 (Fig. 4B-D). Since we anticipated these samples would exhibit staining near the limit of detection of the instrument, a higher concentration of secondary antibody was used to maximize fluorescent signal. Despite this, F12 was the only mAb that produced notable staining intensity, a clear virus population shift, and a higher absolute count of stained virions compared to the isotype control (Fig. 4B-D) within the virus staining gate (Fig. 4B). In contrast, S1-specific F07 and RBD-specific C04 yielded moderate staining intensities (nasal: 80–95 PE MESF; bronchial: 60–105 PE MESF) but labeled relatively few particles compared to F12. These findings suggest that the epitopes targeted by F07 and C04 are less abundant or accessible on viruses produced in ALI cultures. We observed that although all viruses were effectively labeled by mAb F12, differences in the distribution and appearance of virus populations were evident between nasal and bronchial samples. For instance, even within the same donors, both staining intensity and total event counts varied between bronchial and nasal samples, without a consistent pattern across donors.

Across all conditions, the bubble plots (Fig. 4C–D), where bubble size represents particle count and color indicates mean fluorescence intensity (PE MESF), show that intensity and particle abundance do not necessarily correlate. In both nasal (Fig. 4C) and bronchial (Fig. 4D) samples, some antibody–sample combinations exhibit moderate MESF values but low numbers of labeled particles (e.g., C04), whereas others show both moderate intensity and high particle counts (e.g., F12). Notably, F12 consistently labels a larger fraction of virions across donors and tissue types, while C04 and F07 produce measurable fluorescence signals on comparatively few particles. Additionally, variability between nasal and bronchial samples appeared largely donor-dependent rather than systematic, suggesting that the availability of epitopes on S is influenced by both tissue source and host-specific factors. Together, these data highlight the power of flow virometry to resolve heterogeneity at the single-virion level in clinically relevant samples, revealing features of antigen presentation that may be overlooked by bulk techniques.

## DISCUSSION

As biology advances toward increasingly precise single-cell phenotyping, there remains a need for methods that can resolve antigenic heterogeneity at the level of individual virions rather than population-level averages across particle populations^98^. Although traditional flow cytometers often require specialized modifications to detect viral particles^2,16,38^, newer commercially available instruments with enhanced sensitivity^32,99^ now make FV an accessible and practical approach for single-virion analysis.

In this study, we establish FV as a calibrated, single-particle, and high-throughput platform for assessing antigenicity of HCoVs at the single-particle level. While FV has largely been applied to SARS-CoV-2 and MERS-CoV pseudoviruses ^23,55,56,58^, our work extends its use to native virions of HCoVs (229E and OC43) without the need for purification, magnetic enrichment or specialized instrumentation. We show the same FV approach can be applied directly to replication competent SARS-CoV-2, in alignment with similar emerging reports^59^. FV provides a more physiologically relevant view of the viral surface than results generated using recombinant S proteins alone, particularly since virion-to-virion variation in antigenicity may influence infectivity^100^. Accordingly, FV is best viewed as complementary to bulk and structural methods, with particular utility for identifying heterogeneity and prioritizing samples for downstream mechanistic analysis.

Building on this advantage, FV also enables direct comparison of viral populations produced in different cellular contexts. Producer cell type is known to influence virion infectivity as well as its proteomic and surface composition^92,101,102^. Notably, FV has previously been used to highlight viral heterogeneity across cellular models in the context of HIV^36,103^ and to distinguish pleiomorphic influenza virions^104^, underscoring its broader utility for dissecting structural and compositional diversity at the single-particle level. In this study, we demonstrate that FV can effectively resolve cell line-dependent differences in HCoV S antigenicity. Both HCoV-229E and HCoV-OC43 virions propagated in distinct human cell lines exhibited variability, with HCoV-229E showing differences in spike staining intensity and HCoV-OC43 displaying a notably broad side scatter distribution, highlighting the benefit of studying S on virions at the single-particle level to resolve these differences. This variability was particularly pronounced for HCoV-OC43, which displayed a broader range of staining than HCoV-229E—consistent with observations by Schirtzinger et al., who found that HCoV-OC43 propagated in Vero E6, MRC-5, or HCT-8 cells produced different ratios of defective particles^105^. Importantly, we observed that S antigenicity on PV can differ substantially from that on infectious viruses produced in cell lines. While PV models performed comparably for HCoV-229E, they were less representative for HCoV-OC43, indicating that the choice of PV model requires careful validation for each virus system. This discrepancy may also reflect sequence differences between the spike used for pseudovirus production and that of the live virus, as well as additional mutations introduced during passaging of the ATCC-derived live virus stocks, both of which may influence spike antigenicity, particularly within the more variable S1 domain. More broadly, these data emphasize that agreement between PV and native virions cannot always be assumed, even when the same S is being evaluated.

While effective CoV vaccines have been successful utilizing a prefusion-stabilized S antigen approach^62,63^, the extent to which these engineered spikes recapitulate the conformational states and epitope accessibility of S on infectious virions from all CoV genera remains incompletely defined. In our study, staining CoV in the presence of soluble cellular receptors led to measurable changes in antibody staining patterns, consistent with altered epitope accessibility during the transition between pre- and post-fusion states. These findings, which parallel prior observations for the HIV envelope^80,103^, demonstrate FV’s ability to capture structural dynamics that influence antibody binding. Detecting conformational differences in S structure on native virions could provide valuable insight into how receptor engagement and mutation shape antigenicity and may help identify cases in which recombinant immunogens incompletely reflect the antigenic states sampled by infectious particles.

Notably, comparison of FV with ELISA and biolayer interferometry (BLI) revealed broadly consistent antigenic trends for betacoronaviruses using S2 Abs, though discrepancies emerged for 229E S-2P ELISA and FV on native virions. This divergence likely reflects structural differences between recombinant stabilized spikes and those on infectious virions, underscoring the need for cross-platform validation when using recombinant proteins to model native antigenicity. While overall trends were similar, FV’s single-virion resolution provides additional context to interpret these differences and uncover underlying sources of antigenic variation. In this sense, FV can complements bulk assays by revealing whether an apparently intermediate binding signal reflects uniform moderate staining across particles or strong staining of only a virion subpopulation.

ALI cultures are well accepted as a more physiologically relevant model of respiratory virus infections than 2D cultures of epithelial cells^96,106–108^. Strikingly, our results show HCoV-229E virions derived from patient airway ALI cultures displayed reduced labeling compared to those produced in immortalized cell lines, particularly for mAbs targeting the RBD and S1 regions. These findings may suggest that the epitopes targeted by F07 and C04 are less abundant or available on viruses produced in ALI cultures. Interestingly, a similar trend was observed in FV staining of the HIV envelope glycoprotein in samples cultured in cell lines and in peripheral blood mononuclear cells (PBMC)^103^. FV thus provides a unique means to quantify these differences directly and to validate the relevance of cell line–derived viral stocks to human infection. Notably, the ALI experiments represent one of the clearest demonstrations of the value of FV in clinically relevant samples, as bubble-plot analysis showed that fluorescence intensity and stained-particle abundance did not necessarily correlate across donors, tissues, or antibodies. This distinction would be difficult to resolve using bulk measurements alone, particularly in low-signal samples near the limit of detection. This reduced labeling likely reflects physiological differences in S processing or conformation in primary cells, emphasizing the need to consider cellular context when evaluating antigenicity. At the same time, the donor- and cell-dependent differences observed here underscore that virion-associated antigenicity is not fixed but can vary substantially even within a physiologically relevant culture system. Similar cell type- and donor-specific variation in susceptibility has also been reported for Epstein-Barr virus in ALI models^109^, suggesting that such differences are also inherent to this system. Future experiments, assessing antigenicity of HCoVs in patient nasal swabs would provide further detail about viruses which circulate directly in patients, as seen recently for SARS-CoV-2^59^.

Collectively, this work positions FV as a versatile analytical tool for calibrated, single-particle characterization of viral surface proteins. Its capacity to capture particle-level heterogeneity, conformational shifts, and cross-reactive binding patterns complements established biochemical and structural approaches. Importantly, our findings demonstrate that coronavirus S antigenicity is not an intrinsic, fixed property of the virus, but is shaped by the cellular context in which virions are produced, and that commonly used surrogate systems, including pseudoviruses and recombinant spike proteins, do not always fully recapitulate the antigenic landscape of native virions. More specifically, our study shows that FV can be used to compare intact virions across production systems, detect perturbation-dependent changes in antibody binding, and identify clinically relevant heterogeneity in patient-derived airway cultures without prior particle enrichment. Looking forward, FV could be applied to other enveloped viruses to assess vaccine integrity, monitor antigenic drift, or evaluate therapeutic antibody binding in physiologically relevant contexts. When used alongside orthogonal structural, biochemical, and sorting-based approaches, FV provides a practical framework for defining how viral antigenicity varies across particles, samples, and production systems in ways that are obscured by bulk measurements.

### Limitations of the study

Although indirect staining with full length Abs successfully labeled virions in this study, future use of nanobodies may reduce steric hindrance and improve access to concealed epitopes. While direct staining was less effective here than indirect staining, further optimization of conjugation efficiency may enhance signal detection, particularly for low-abundance antigens. The cytometer used here is designed for particles ≥80 nm, potentially limiting the detection of biological particles below this threshold. While our FV assay allows for every particle in the preparation to be viewed —avoiding selection bias introduced by the requirement of magnetic nanoparticles (MNPs) or fluorescent triggering using nucleic acid dyes—this approach increases the importance of antibody titration and inclusion of appropriate controls to distinguish specific signal from background fluorescence. Fluorescence calibration was performed using beads designed for cellular flow cytometry; as a result, some quantitative values may be less accurate due to use near the lower limit of the standard curve. The field would benefit from expanded availability of small particle calibration standards to improve quantitation and reproducibility. Lastly, EVs were not analyzed in this study. Distinguishing EVs from viruses remains an important challenge^73^, and additional work is required to resolve these differences using FV and complementary antibody-based approaches. Future work will incorporate newer models of cytometers that are dedicated to small particles^99^ and virus sorting which remains an important complementary technique that can advance flow virometry studies^1,16,17,24,110^. It should be noted that the HCoV-229E and HCoV-OC43 strains used here are tissue culture adapted, which may introduce antigenic differences in spike compared to currently circulating clinical isolates. Future studies should aim to characterize spike antigenicity in primary isolates to better define these effects in a clinically relevant context.

## Author Contributions

JB and KSC conceptualized the project. JB, CO, CD, CH, GO, LE, TM, VL performed all experiments and data analysis. CD, HM, JJ, SM, NST, LHT, DH provided key reagents (i.e., plasmids, virus stocks, antibodies). JB, NAC, SW, PK, KSC provided supervision and funding. JB wrote the original manuscript draft. JB and KSC revised and polished the manuscript with input from all coauthors. All authors have read and agreed to the published version of the manuscript

## Acknowledgements

We thank the staff at the Center for Extracellular Vesicle Research at the Beth Israel Deaconess Medical Center, especially Garrett Haskett and Vanessa Costa for help with sample acquisition. We thank Fidan Baycora for administrative support and Emily Hobbs for grant support, respectively. We would like to thank Dr. Sunil Singhal and Azra Din for their help in obtaining the paired nasal and bronchial samples for establishing primary epithelial cultures. All illustrations were created with BioRender. The following reagents were obtained through BEI Resources, NIAID, NIH: Homo sapiens Lung Carcinoma Cells (A549), Serum-Free, (NR-52268) Human Coronavirus OC43 in HRT-18G Cells (NR-56241), Human Coronavirus 229E (NR-52726), and Monoclonal Anti-SARS-CoV-2 S RBD Chimeric Antibody (NR-55408), HIV-1 SG3ΔEnv (ARP-11051), contributed by Dr. John C. Kappes

## Declaration of generative AI and AI-assisted technologies

During the preparation of this work, the first-author (JB) used ChatGPT for writing revisions to improve the readability and language of the manuscript. After using this tool or service, the authors reviewed and edited the content as needed and take full responsibility for the content of the publication.

## Data and code availability

Source data supporting the findings of this study are available within this paper and its supplementary files. Source data have also been deposited to Figshare and can be accessed at DOI: 10.6084/m9.figshare.32254299. Additional source data inquiries should be directed to the corresponding author, Kizzmekia Corbett-Helaire.

## Material and resource availability

All requests for reagents and resources should be directed to the corresponding author, Kizzmekia Corbett-Helaire and will be made available after completion of a Material Transfer Agreement (MTA). If the material was obtained under use restriction, the inquiry will be forwarded to the appropriate party.

## Method Details

### Cell culture

All cells were grown in a 5% CO_2_ humidified incubator at 37 °C in complete media containing 10% FBS (GeminiBio, Cat# 100-106) and 1% penicillin/streptomycin (Life Technologies, Cat# 15140122) unless otherwise noted. A549 (BEI Resources, Cat #NR-52268), Vero E6 cells (from Dr. Phyllis Kanki) and Huh7.5 cells^111^ (provided by Deborah R. Taylor, US Food and Drug Administration, RRID: CVCL_7927) were cultured in Dulbecco’s modified Eagle’s medium (DMEM, Gibco, Cat# 10313-021). MRC-5 cells were cultured in EMEM (ATCC, Cat# 30-2003) supplemented with Antibiotic-Antimycotic (Gibco, Cat# 15240062). HCT-8 (HRT-18) cells (ATCC, Cat# CCL-244) were cultured in RPMI (Gibco, Cat#11875-093).

#### Primary human nasal and bronchial cell cultures

Donor matched sinonasal and bronchial cells were obtained from patient donors with informed consent, per protocol approved by the University of Pennsylvania Institutional Review Board (protocol #813004, PI Sunil Singhal). A detailed protocol for culturing primary cells at ALI was described previously^93^. Briefly, specimens were grown in PneumaCult™-ALI Medium (STEMCELL Technologies 05001) supplemented with heparin (STEMCELLl Technologies 07980) and hydrocortisone. Once cells reached 80% confluency in 0.4 µM pore transwell inserts, the apical growth media was removed, and the basal differentiation media was replaced every 3-4 days for a minimum of 4 weeks. Prior to infection epithelial morphology and cilia beating were confirmed by microscopy.

### Flow virometry

Flow virometry was performed using a Beckman Coulter CytoFLEX S with a standard optical configuration. All samples were acquired for 15-30 seconds at a sample flow rate of 10 μL/min using the tube or plate loader. For all labeling experiments, cell-free supernatants containing virus were stained at their undiluted titer. For indirect labeling, viruses were incubated with of unlabeled primary mAbs (0.2-1 µg/mL) overnight at 4 °C, followed by a three-hour incubation with 0.2-1.2 µg/mL of PE labeled secondary antibody (anti-human PE BioLegend, Cat# 410708; anti-mouse PE, Invitrogen, Cat# A10543). For direct labeling, the CTD antibody H501-0008 was conjugated using the LYXN Rapid RPE antibody conjugation kit (Bio-Rad; Cat # LNK022RPE). Serial dilutions of the conjugated Ab were used to empirically determine the optimal staining concentration. Similar staining procedures as described above were used. For staining experiments with recombinant proteins (DPP4, BioLegend, Cat# 764106; APN, SinoBiological, Cat# 10051-H08H), virus was incubated with 5 µg/mL final of protein for 20 min before staining was completed as described above. After staining, viruses were fixed in a final concentration of 2% PFA for 20 min before being diluted with PBS (ThermoFisher Scientific, Cat#10010023) for acquisition. BD Quantibrite PE beads (Cat# 340495, lot# 75221) were used for fluorescence calibration while NIST traceable size standards (Thermo Fisher Scientific) were used for light scatter calibration. Calibration was performed using FCM_PASS_ software (https://nano.ccr.cancer.gov/fcmpass) as previously described^33,112^. Additional details on calibration, antibody titrations and MIFlowCyt-EV framework^113^ controls are described in File S1.

### Plasmid Design and Construction

For recombinant S protein production, codon-optimized S sequence of SARS-CoV-2 Wuhan (GenBank accession QHD43416.1), HKU1 (GenBank accession, ABC70719.1), OC43 (GenBank accession, AIL49484.1), 229E (GenBank accession, NP073551.1), NL63 (GenBank accession, YP 003767.1), MERS (GenBank accession, AFY13307.1) previously reported in this study^114^, were obtained from the Vaccine Research Centre (VRC). All S constructs contain a C-terminal foldon trimerization motif (YIPEAPRDGQAYVRKDGEWVLLSTFL) and an octa-histidine tag (HHHHHHHH) for purification. A human rhinovirus 3C protease recognition site (GSRSLEVLFQGP) was inserted to enable tag cleavage after purification. In addition, the S proteins were prefusion stabilized via two proline substitutions and contain S1/S2 furin cleavage site modification^60,76^. For antibody production, previously designed plasmids encoding heavy and light chains of monoclonal antibody, H501-008, H501-101, D12, F11, G2, IgG22, G4; and fp.006 were provided by VRC^60,115^ and Davide F. Robbiani lab at Institute for Research in Biomedicine^82^. Plasmids encoding heavy and light chains of mAbs CC40.8 and 54043-5 were obtained from Andrabi lab (Scripps Research Institute)^90^ and Georgiev lab (Vanderbilt Vaccine Center)^84^. Plasmids expressing heavy and light chains of mAbs (C04, F07, F12, S2P6, 76E1) were synthesized and cloned into pcDNA 3.1(+) mammalian expression vector by GenScript.

### Protein Expression and Purification

S proteins were produced by transiently transfecting Expi293F cells with the plasmid using polyethyleneimine “MAX” (PEI-MAX, polyscience, Cat# 24765). Cell culture supernatants were harvested on day 5 post-expression. Secreted S proteins were purified by StrepTrap HP affinity column chromatography (Cytiva). Monoclonal antibodies were produced by transiently co-transfecting Expi293F cells with heavy and light chain plasmids using polyethyleneimine in a 1:1 ratio. Cell supernatants were collected 6 days post-transfection, and antibodies were purified with protein A agarose (Cytiva). Bound mAbs were eluted with 100 mM glycine at pH 2.7 into 1/10th volume of 1 M Tris-HCl pH 8.0, buffer exchanged into 1x PBS, pH 7.4 and flash-frozen for long-term storage at -80 °C. Both purified spikes and mAbs were validated for binding by antigenicity ELISA before long-term storage and use.

### Pseudovirus production

MERS-CoV pseudoviruses were produced through transfection of HEK293T cells using Polyjet In Vitro Transfection Reagent (SignaGen Laboratories, Cat# SL100688). Pseudoviruses were generated using 1 μg of SG3ΔEnv plasmid (ARP-11051) or VRC5602^116^ as the viral backbone, and 0.5 μg of the MERS-CoV S protein. Viruses were harvested 48 hours after transfection. HCoV-229E pseudoviruses were produced as previously described^117^. VSV-based HCoV-OC43 pseudoviruses were produced as described by McMcCallum et al.^118^. HCoV-OC43 and HCoV-229E S plasmids were cloned in house into a eukaryotic expression vector pCDNA3.1 (GenScript) with the full-length S coding sequences lacking the ER retention signal. SARS-CoV-2 and MERS-CoV S plasmids were obtained from Addgene (Cat# 155297^119 &^ 170448 respectively).

### Cell line-derived virus production

HCoV-OC43 (BEI, Cat# NR-56241) was used for infection in the human cell lines MRC-5, Huh7.5^111^, HCT-8, and A549 while HCoV-229E (BEI, Cat# NR-52726) was used for infection in the MRC-5, Huh7.5 and HCT-8 cell lines. Viruses were passaged 2-3 times in each respective cell line before use in experiments. Cells were infected at 80% confluence with virus isolates until the time of harvest 4-5 days after infection. Fresh media was added to the cells as needed before time of harvest. Cell culture supernatants containing virus were centrifuged at 300 x g for 5 minutes to remove cellular debris before being aliquoted and stored as unfiltered viral stocks at −80 °C until use in subsequent assays.

#### ALI virus production, replication curves, and plaque assays

Infections of primary nasal cultures at air-liquid-interface was previous described^93^. Nasal and bronchial ALI cultures were infected on the apical surface with HCoV-229E at MOI 0.1 PFU/mL in serum-free DMEM for 1 hour at 33°C, followed by removal of virus inoculum and three PBS washes. Apically shed virus was collected at 96 hours post infection in PBS. Infectious particles were quantified by plaque assay on Huh7 cells.

### Biolayer Interferometry

Binding kinetics of CoV S proteins to mAbs were carried out using an Octet RH96 system (Sartorius) at 30 °C, shaking at 1,000 rpm. Monoclonal antibodies were diluted to 40 - 50 µg/mL in kinetic buffer (1x HBS-EP + buffer, cytiva #BR100669). S Proteins were diluted to 100 µg/mL in kinetic buffer and serially diluted two-fold for a final concentration of 6.25 µg/mL. Reagents were added to a solid black, tilted-bottom 384-well plate (Geiger Bio-One) at 90 µL per well. Following manufacturer’s instructions, mAbs were immobilized onto protein A Fc capture probes (Octet ProA Biosensors, Cat #18-5010) for 180 s to capture levels of 1−1.5 nm. Biosensor tips were then equilibrated for 180 s in kinetic buffer before binding assessments of CoV S proteins for 300 s of association, followed by dissociation for 120 s. Data analysis and curve fitting were done with the Octet software, v 8.0. The data were baseline-subtracted before exporting and plotting in GraphPad Prism. Plots show association and dissociation steps. Experimental data were fitted with a 1:1 binding model in the Octet software, v 8.0.

### ELISA

96-well Nunc MaxiSorp flat-bottom plates (ThermoFisher, Cat# 44-2401-21) were coated with 1 µg/mL of recombinant S-2P CoV S proteins diluted in PBS for an overnight incubation. Plates were subsequently washed three times following every step using PBS + 0.05% Tween 20. Blocking was performed with 5% nonfat Difco skim milk (BD, Cat# 232100) for one hour. Anti-S mAbs were diluted at 2 µg/mL in blocking buffer and were incubated for 1 hour before washing and the addition of Goat anti-Human secondary antibody (Fortis Life Sciences, Cat# A80-119P). 1-Step TMB ELISA Substrate (Thermo Scientific, Cat#34029) was added to wells before absorbance 405 nm was read on a SpectraMax iD5 Multi-Mode Microplate Reader (Molecular Devices).

### Data analysis

Data visualization and statistical analyses were performed with GraphPad Prism v10. All flow virometry data were analyzed using FlowJo software version 10.10.0. Bubble plots were constructed in R (v4.4.0) using the *ggplot2* package. For each antibody–virus combination, the mean PE MESF (staining intensity) and event count obtained from the virus gate were plotted, with color indicating staining intensity and bubble size proportional to event count.

