## Supplemental figures for "Evaluating Spike Antigenicity across Endemic Human Coronavirus Models using Flow Virometry"

### Fig S1. Workflow and validation of an FV platform for quantifying HCoV-229E S antigenicity

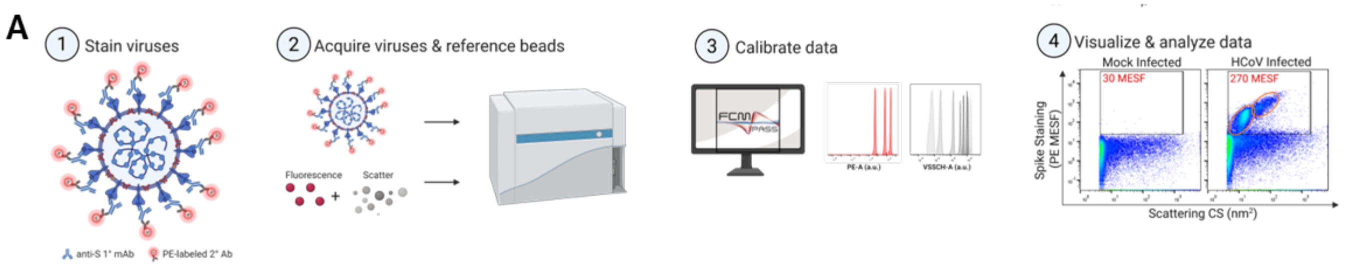


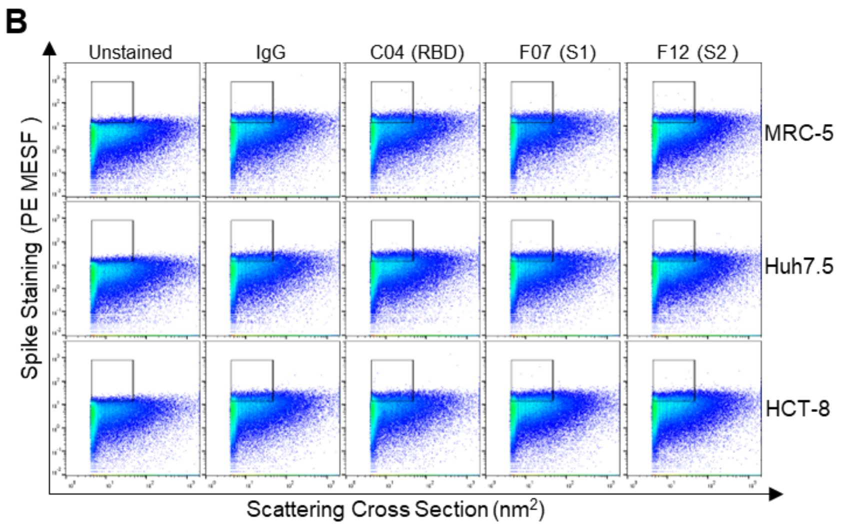


C

| Producer Cell | Stock | TCID50 |
|---|---|---|
| MRC-5 | 1 | 5.82E+07 |
| | 2 | 1.55E+06 |
| Huh7.5 | 1 | 3.89E+07 |
| | 2 | 3.89E+07 |
| HCT-8 | 1 | 2.82E+04 |
| | 2 | 1.88E+04 |

**Figure S1. Workflow and validation of an FV platform for quantifying HCoV-229E S antigenicity.** (A) General workflow for indirect staining flow virometry assays. This protocol served as the basis for all staining procedures. (1) Viruses in the culture supernatants of infected cells were stained with monoclonal antibodies (mAbs) targeting different CoV S protein epitopes, followed by a PE-conjugated secondary antibody. (2) NIST-traceable sizing beads and BD Quantibrite fluorescence reference beads were used for scatter and fluorescence calibration respectively, enabling reporting in standardized units. Additional assay controls were acquired as outlined in the MIFlowCyt-EV framework. All samples were acquired on a CytoFLEX S cytometer at the lowest sampling rate. (3) Data calibration was performed using FCMPASS and (4) calibrated data were analyzed using FlowJo. Orange cirlces denote two distinct mAb stained populations. (B) Matched mock-infected supernatants from the human cell lines MRC-5, Huh7.5, and HCT-8 (top to bottom rows) were stained with S mAbs targeting the HCoV-229E RBD (C04), S1 (F07) or S2 (F12) domains, or an isotype control antibody (IgG), followed by a PE-labeled secondary antibody. Gates for positive staining were set above the background fluorescence of the isotype control (>15 PE MESF on the y-axis). (C) TCID50 data from 229E virus stocks used in experiments.

### Fig S2. Establishing an FV platform to assess S antigenicity on HCoV-OC43

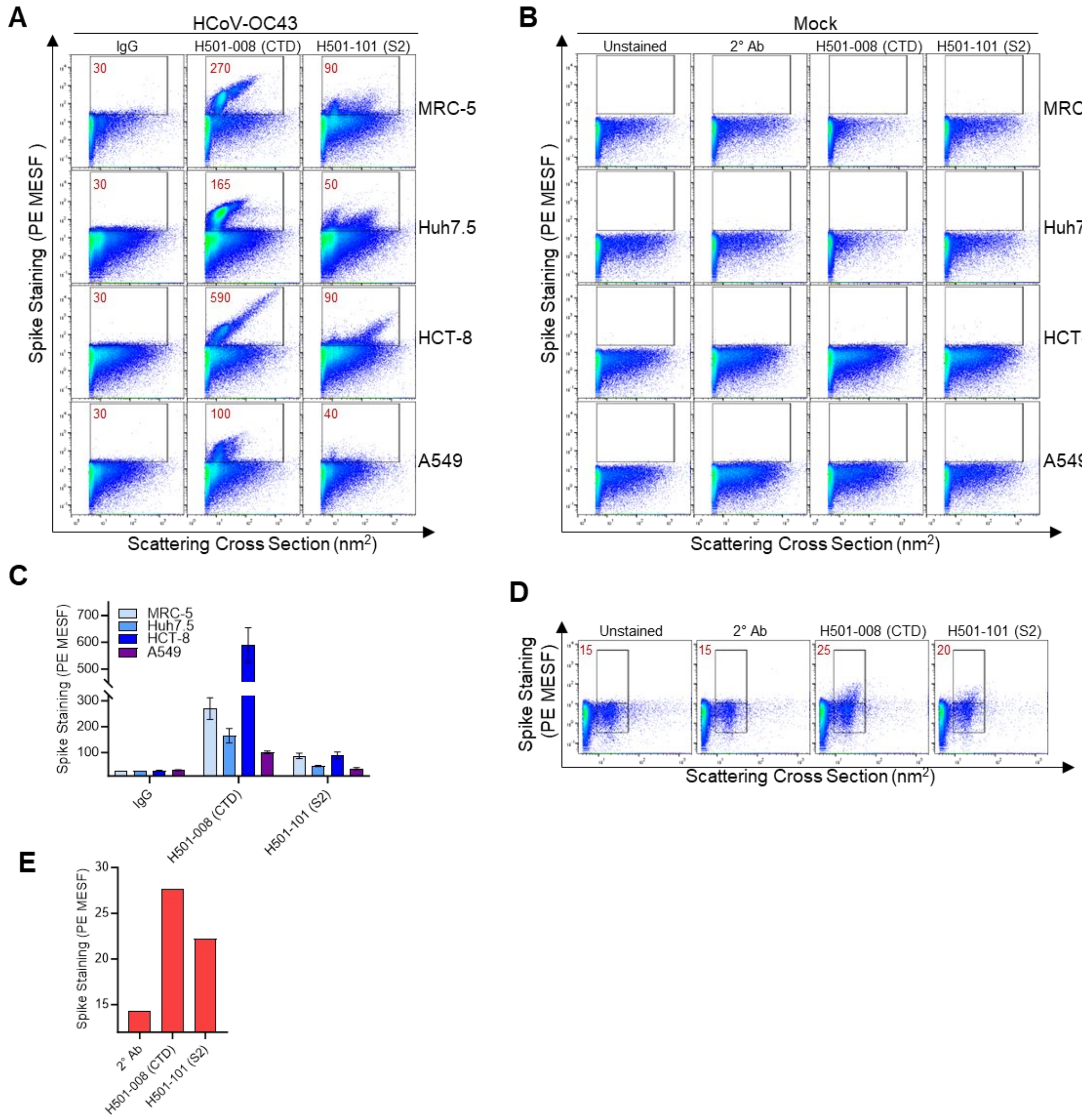


**Figure S2. Establishing an FV platform to assess S antigenicity on HCoV-OC43.** (A) Supernatants from HCoV-OC43-infected human cell lines MRC-5, Huh7.5, HCT-8 and A549 (top to bottom rows) were stained with S mAbs targeting the CTD (H501-008) or S2 (H501-101) domains, or an isotype control antibody (IgG), followed by a PE-labeled secondary antibody. Gates for positive staining were set above the background fluorescence of the isotype control (>25 PE MESF on the y-axis). Calibrated mean PE fluorescence values derived from the gated regions are shown in red. (B) Mock-infected supernatants from the human cell lines MRC-5, Huh7.5, HCT-8 and A549 (top to bottom rows) were stained as in Fig. 1. Gates for positive staining were set above the background seen from secondary antibody alone (2° Ab). (C) Quantitative comparison of staining results from the gates in (A), with bars representing mean PE MESF values ± standard deviation, wherein light blue = MRC-5, blue = Huh7.5, dark blue = HCT-8, and purple = A549. Results are representative of at least three replicates**.** (D) OC43 pseudoviruses produced by transfection of 293T cells were stained directly in cell culture supernatants using S mAbs targeting the CTD (H501-008) or S2 (H501-101) domains, followed by a PE-labeled secondary antibody. Gates for positive staining were set above the background seen from secondary antibody alone (2° Ab). Calibrated mean PE fluorescence values (MESF) from gated regions are shown in red. (E) Quantitative comparison of staining results from the gates in (D).

### Fig S3. Comparison of spike labelling using direct and indirect staining

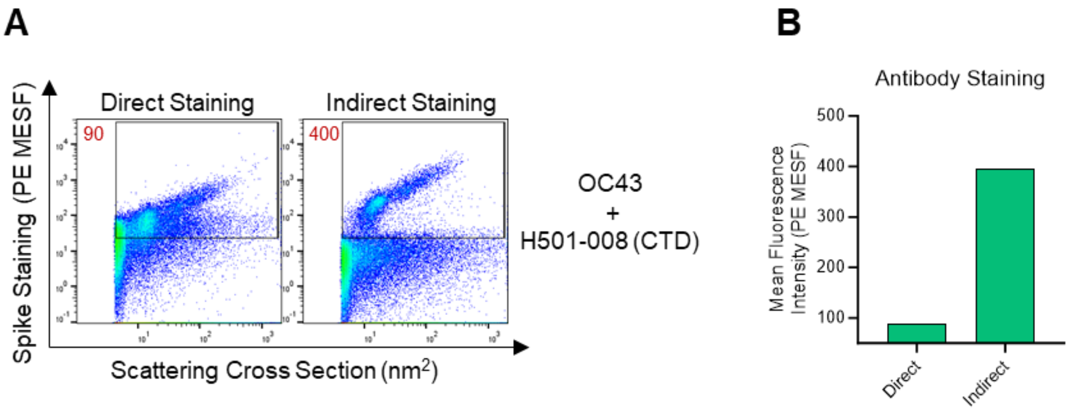


**Figure S3. Comparison of spike labelling using direct and indirect staining.** (A) OC43 grown in MRC-5 cells was stained with a CTD-specific antibody (H501-008) using either an unlabeled primary antibody (indirect) with a PE-conjugated secondary antibody or an in-house PE-conjugated primary antibody (direct). Gates were set based on the positive signal observed in the indirect staining plots. Calibrated mean PE fluorescence values derived from the gated regions are shown in red. (B) Quantitative comparison of staining results from the gate in (A).

### Fig S4. Assessing spike epitope availability in the presence of soluble cellular receptor, DPP4

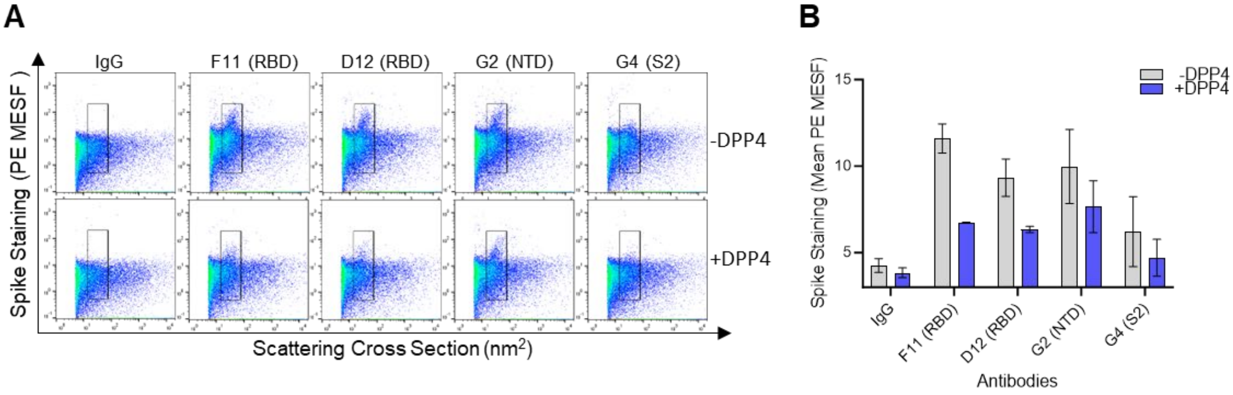


**Figure S4. Assessing spike epitope availability in the presence of soluble cellular receptor, DPP4.** (A) Flow virometry dot plots showing MERS-CoV pseudovirus staining in the presence (bottom row) or absence (top row) of soluble DPP4. Gates are drawn on the total virus population to allow for modest changes in staining intensity. (B) Quantification of staining in (A), showing mean PE MESF ± SD from the gated regions wherein gray = absence of DPP4 and blue = presence of DPP4. Results are shown from one experiment tested in duplicate and are representative of three experimental replicates.

**Fig S5. Comparing S2 antibody binding against HCoV in BLI (1/2)**

**A** 229E

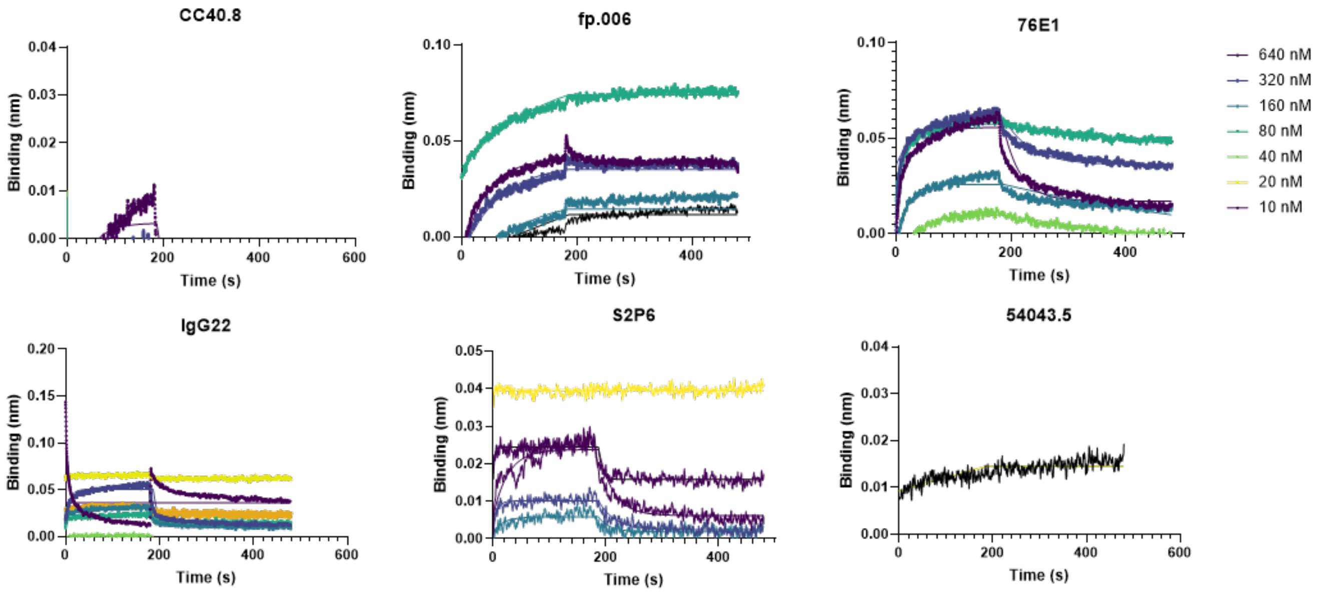


**B** OC43

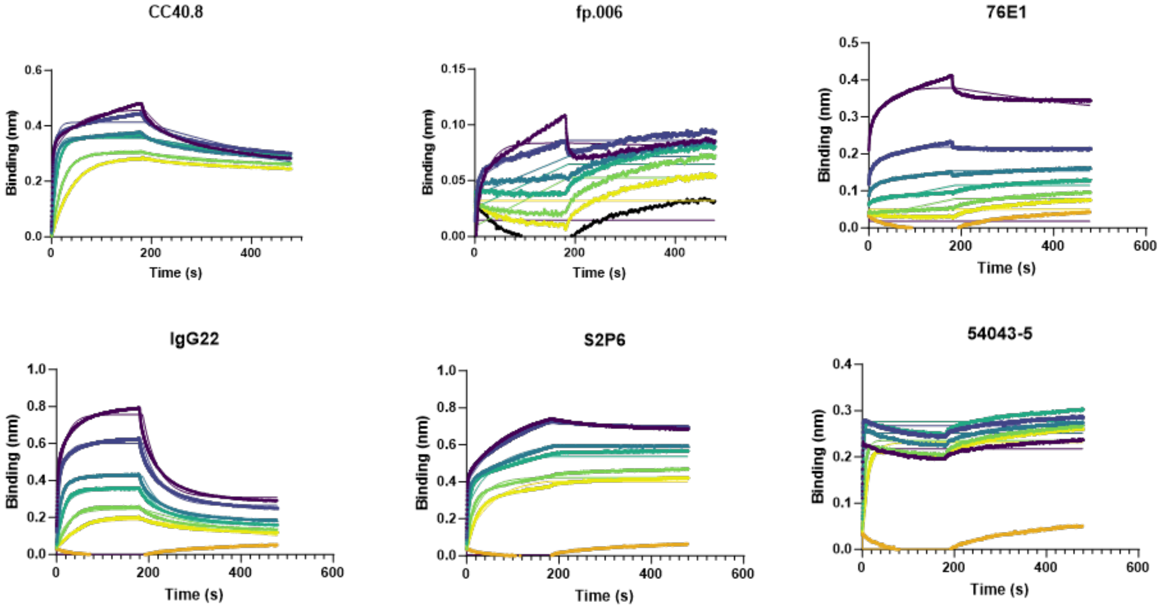

**Fig S5. Comparing S2 antibody binding against HCoV S-2P in BLI (2/2)**

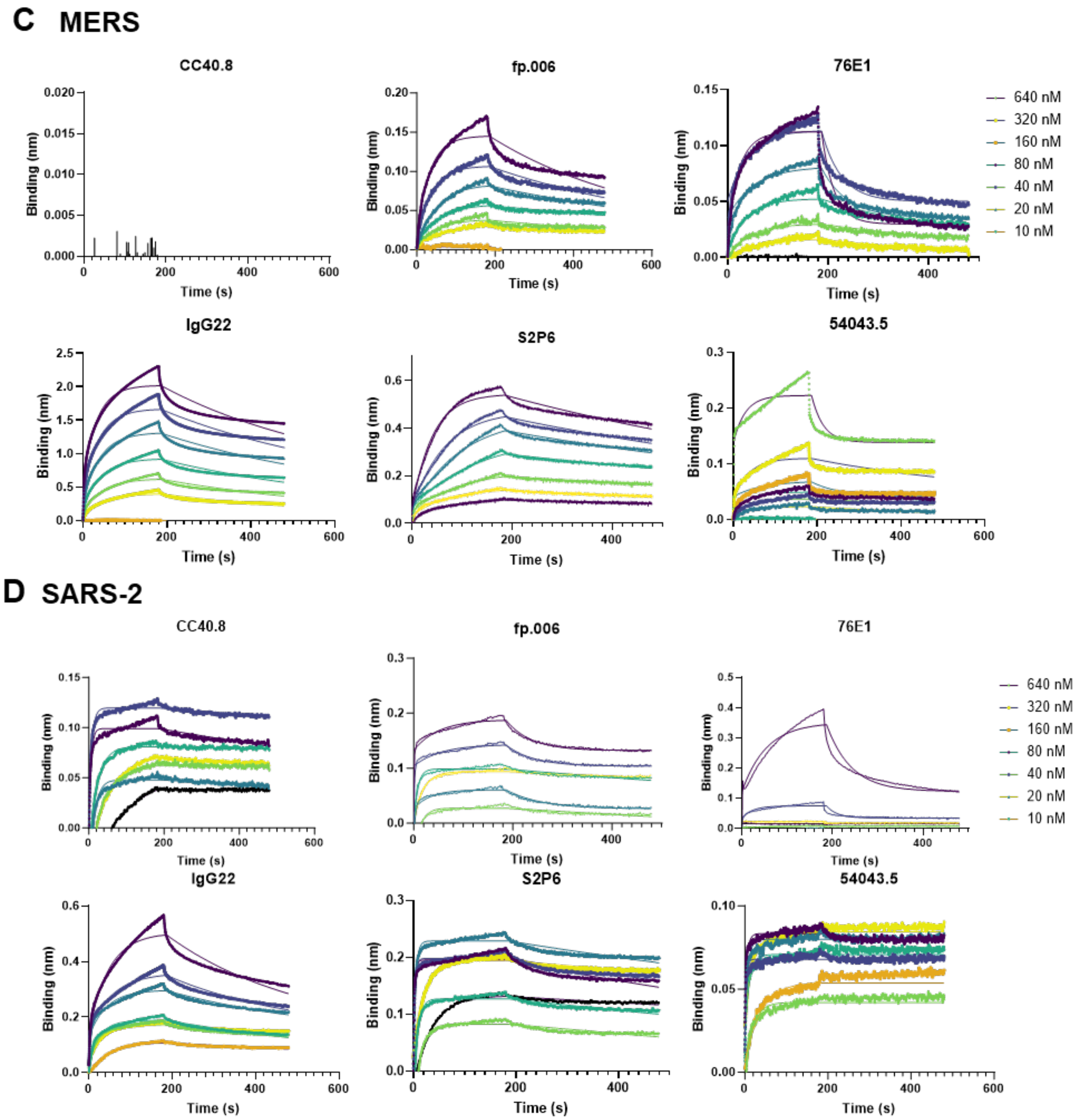


**Figure S5. Binding affinity of recombinant coronavirus spike (S-2P) proteins to immobilized S2 antibodies measured using BLI.** Representative binding curves are shown for (A) HCoV-229E, (B) HCoV-OC43, (C) MERS-CoV, and (D) SARS-CoV-2 spike proteins. Each sensogram depicts real-time association and dissociation kinetics obtained using serial dilutions of the S-2P proteins. Different colored traces denote the concentrations used for each condition. Data are representative of two biological replicates using distinct batches of proteins and antibodies.
